# Horizontal Gene Transfer of *E. coli* by direct DNA uptake from extracellular liquid phase

**DOI:** 10.1101/2025.11.21.689801

**Authors:** Curtis Wood, Zachary Smith, Lauren Johnson, James Weifu Lee

**Affiliations:** Old Dominion University, Department of Chemistry & Biochemistry, Norfolk, 23529, U.S.A

**Keywords:** Horizontal gene transfer, genetic transformation, Bacteria E. coli, direct uptake of DNA from extracellular liquid, biosafety research, environmental DNA

## Abstract

The “fate of naked DNA” or “environmental DNA (eDNA)” from genetically engineered (GE) biomasses in relation to possible horizontal gene transfer (HGT) is an emerging area of biosafety research. Using a broad-host plasmid (pAM5409) as a mimic of eDNA containing a yellow fluorescent protein gene (*yfp)* and conferring spectinomycin/streptomycin (Sp/Sm) resistance, we incubated each of the *E. coli* strains DH5α, DH10B and K12 with the plasmid eDNA to evaluate HGT through direct uptake of plasmid DNA from extracellular liquid phase. According to experimental observations, the HGT can occur within 1 day of donor (eDNA) and recipient cells liquid incubation in *E. coli* strains DH5α and DH10B, but not in K12 which has its intact genes encoding restriction enzymes and endonucleases. HGT occurrences were observed by analyzing colony forming units (CFU) expression of yellow fluorescent protein (YFP). The HGT frequency by direct uptake of plasmid DNA from the liquid phase was determined to be 1.2 x 10^-8^ per cell day for both DH5α and DH10B. The frequency of HGT was slightly higher than that of spontaneous mutation for natural antibiotic resistance development that was estimated to be 4.8 x 10^-9^ per cell day. Extension of the liquid eDNA and recipient cells incubation time from 1 day to 3 and 5 days did not increase the HGT frequency under the current experimental conditions. Despite K12 failing to form CFUs following exposure to eDNA, this does not imply K12 did not uptake eDNA; eDNA acquired could have been fragmented by K12 restriction enzymes into non-functional oligomers incapable of HGT.

## Introduction

Horizontal gene transfer (HGT), the acquisition and expression of exogenous DNA by an organism, is a well-established phenomenon in prokaryotes that can occur by one of three mechanisms: genetic transduction, conjugation or transformation.^1-4^ Genetic transduction ^5^, the introduction of viral DNA containing mobile genetic elements (MGEs) into bacteria via infectious bacteriophages, can result in a dormant viral state where the MGE is incorporated into the bacterial chromosome, also known as a prophage. Upon induction, the prophage sequence is replicated out or looped out (via homologous recombination) of the host chromosome, forming a functional plasmid containing the genes for synthesis of new phages. Depending on the prophage nature, some of the host chromosome downstream from the prophage sequence is also replicated out and packaged into newly synthesized phages with the rest of the MGE sequence. Upon infection of a new bacteria, the phage again introduces its viral sequence but also the sequence from the previously infected bacteria, potentially transferring in functional genes.^1^ Conjugation is another HGT process that occurs between two bacteria, one acting as the MGE donor and the other as the MGE recipient. The donor utilizes a transmembrane pilus conduit to physically dock with the recipient bacteria before exporting an MGE through the pilus into the recipient bacteria. ^6^ The pilus and exporting machinery differ between bacteria but utilize homologous structures and genes which have been classified as type 4 secretion systems (T4SS) which require the donor cell have a suite of T4SS genes, an origin of transfer (OriT) upstream of the MGE to transfer and a *mob* gene encoding an OriT recognizing relaxase ^7^. The induction of conjugation results in the translation and installation of the coupling proteins and membrane-bound pilus which can attach to a recipient bacteria, expression of the *mob* gene protein which binds the OriT, removes a single strand of the MGE and shuttles it to the T4SS for export through the pilus.^8^ Both the transduction process and the conjugation mechanism continue to be topics of molecular biology research today while the third, bacterial transformation, garners much less attention despite being discovered first and routinely exploited in synthetic biology labs.

Genetic transformation loosely refers to the cellular uptake and utilization of exogenous DNA acquired during a cellular state known as competency. The artificial induction of cellular competency via calcium has been the workhorse of molecular cloning since the 1970’s when the *E. coli* K12 strain was first transformed ^9^. Since then only approximately 80 bacterial species have been shown to possess a naturally competent state, the vast majority of which are pathogenic strains as they are the most studied. ^10^ A host of genes and their homologues responsible for both the regulation of competency (*recA, ssb* and the *com* operon) and DNA import machinery (the *pil* and *fim* operons) have been identified despite vast differences in the environmental conditions invoking competence ^11^; *H. pylori* constitutively expresses DNA import machinery selective for DNA originating from other *H. pylori* bacteria while *S. pneumoniae* becomes competent for any DNA via quorum sensing ^3^. Interestingly, *E. coli*, despite being engineered into multiple strains capable of induced transformation as well as possessing genes homologous to those responsible for natural competence in other bacteria, was thought to not have a naturally competent state until recently.^12^ Some of the genes mutated to confer MGE stability in engineered *E. coli* K12 strains (**Table 1**) are similar to those involved in regulating competency in other bacterial species ^13^. K12 (a.k.a. MG1655) is the most genetically similar to wild-type *E. coli*, only lacking the fertility F plasmid and bacteriophage λ MGEs, while DH5α and DH10B are K12 derivatives engineered to be ideal plasmid factories and repositories; genes encoding restriction enzymes, endonucleases and recombinase have been mutated or deleted, crippling the cell’s ability to degrade foreign DNA, preform homologous recombination and increase mutational rates in response to SOS activation, which in turn increases plasmid retention and stability. Given that *E. coli* is found ubiquitously in soil and freshwater sources as a saprotrophic bacteria, in mammalian, avian & reptilian digestive tracks as both a commensal and pathogenic resident and in laboratories as a chassis for genetic engineering, a better understanding of *E. coli* and other soil bacteria’s capacity for HGT utilizing environmental DNA (eDNA) is warranted in the context of biosafety research.

**Table 1.**
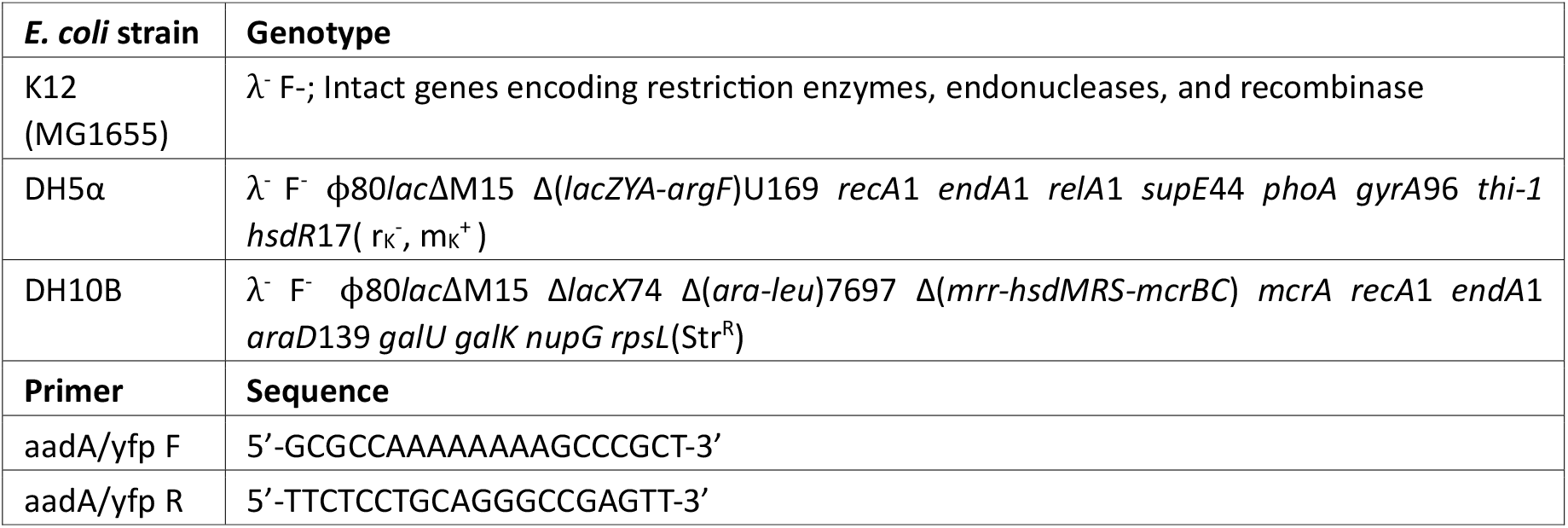
*E. coli* strain genotypes and pAM5409 primer sequences utilized.

The acquisition of eDNA from genetically engineered (GE) biomasses ^14-16^ such as GE crop composts and farmed animals fed GE forages ^17^ has become an emerging area of biosafety research; studies have demonstrated eDNA can remain in soils for as long as 8 years ^18^ and that HGT of MGEs conferring antibiotic resistance from *E. coli* to bacteria in manure amended soil can occur with a probability of 10^-5^ – 10^-4^ recipients per donor.^19^ Previous research in our lab demonstrated HGT of MGEs conferring kanamycin resistance to *E. coli* DH5α that was co-cultured with the GE thermophilic cyanobacteria *T. elongatus* BP1 harboring a pUC based plasmid containing the antibiotic kanamycin resistance MGE.^20,21^ Industrially cultured GE cyanobacteria represent another potential reservoir of antibiotic resistance MGEs ^16^. Independent studies ^22-25^ have applied synthetic biology in cyanobacteria, such as *Synechococcus elongatus* PCC7942 and *Anabaena* (*Nostoc*) PCC 7120, for photobiological synthesis of biofuels and bioproducts such as isobutanol, 1-butanol, columbamide ^24^ and cryptomaldamide ^25^. To efficiently transfer MGEs designed and maintained in *E. coli* to cyanobacteria, vectors capable of replication in multiple bacterial hosts are required, such as the recently developed broad-host-range plasmid pAM5409, a RSF1010 derivative, used to create GE cyanobacteria for bioprospecting.^26,27^

Plasmid pAM5409 contains an origin of replication OriV, an origin of transfer OriT (RK2BOM), the basis of mobility genes *mobA(Y25F), mobB, mobC* as well as their promotors (P1-4) and the suite of *rep* genes (A,B,C,E & F) from pRL1383a, an RSF1010 plasmid derivative. Additionally, pAM5409 harbors *aadA* conferring resistance to spectinomycin/streptomycin (Sp/Sm) and the yellow fluorescent reporter *yfp* (**Figure 1**). The OriT and *mob* genes ensure the plasmid is mobilizable via conjugation should the host contain compatible T4SS genes while the OriV and *rep* genes ensure the plasmid replicates independent of the host’s DNA polymerase affinity for the OriV. While these traits make pAM5409 an attractive vector for genetic engineering, they are the same traits that make it potentially promiscuous within a microbiome and hence our choice to utilize pAM5409 as an example to mimic a source of eDNA.

**Figure 1.**
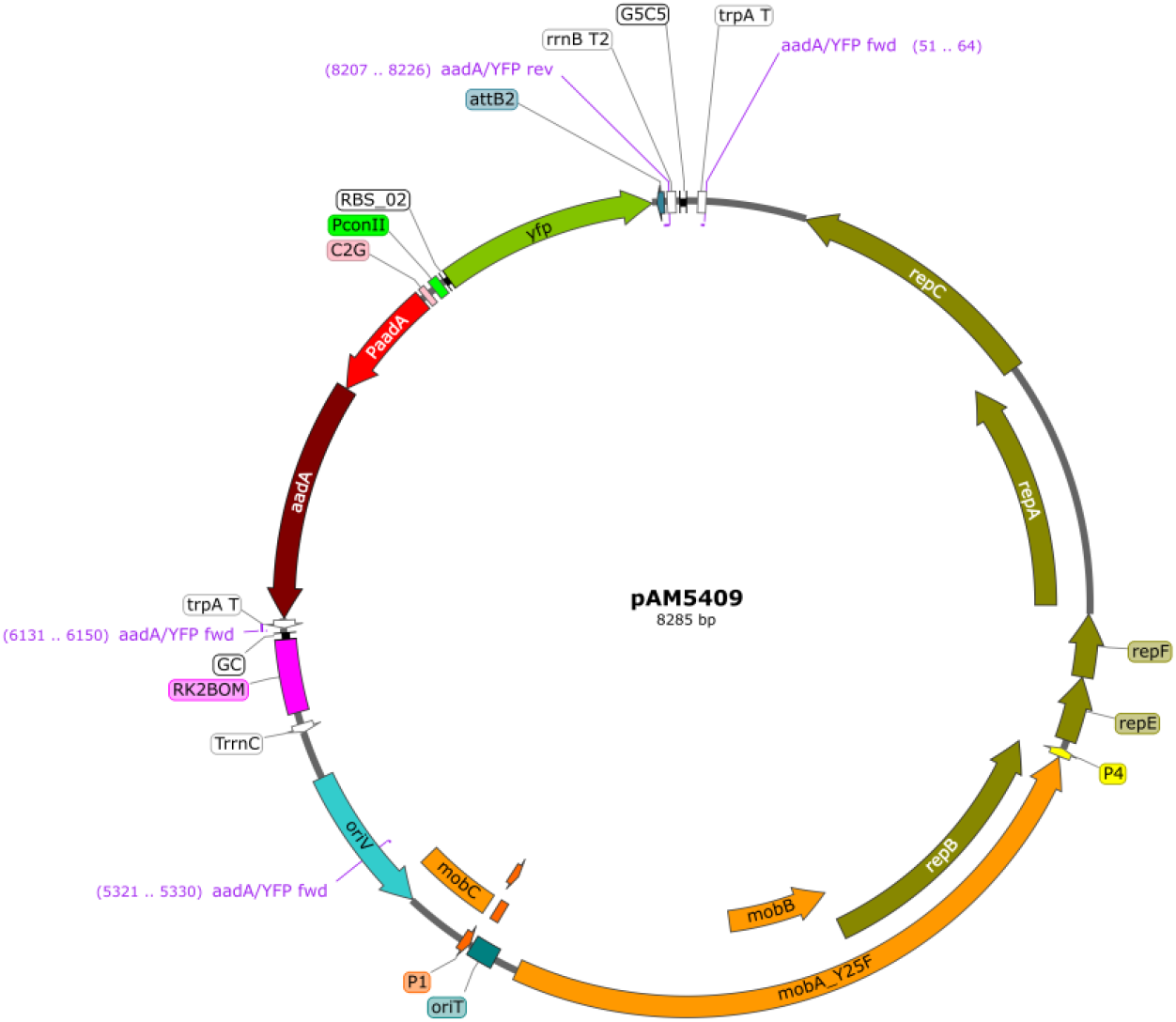
Map of plasmid pAM5409 colored by the following scheme: mobilizing system = orange, replication system = mustard, fluorescent reporter = green, origin of replications = blue, antibiotic resistance = red, termination sequences = white, primer targets = purple.

In this article, we will report our latest experimental study that employed plasmid pAM5409 as a mimic of eDNA to determine the HGT probability to *E. coli* K12, DH5α, and DH10B strains by their direct uptake of plasmid DNA from extracellular liquid phase. As environmental bacteria would be subject to a limited organic nutrient environment, we chose to use a minimal medium comprised of BG-11, a freshwater analogue used to photoautotrophically culture cyanobacteria, mixed with conventional Luria Bertani medium (LB) in an 80:20 ratio as the extracellular liquid phase medium. Plasmid pAM5409, the HGT vector, contains an MGE conferring Sp/Sm resistance as well as a yellow fluorescent reporter gene which is used to display the successful transformation of *E. coli*. We also examined if exposure to a sublethal concentration of antibiotics Sp/Sm influences the probability of HGT; previous studies have established that sublethal antibiotic exposure can lead to an increased number of antibiotic resistant bacteria in a population through a variety of mechanisms.^28,29^

## Results

### Colony-forming unit assays and verification of HGT events by fluorescent imaging analysis of colony YFP expression

In the experiments as illustrated in Figure 2 and detailed in the Methods and Materials section, each of the three *E. coli* strains (K12, DH5α and DH10B) was cultured in 30 ml of LB liquid medium in a flask placed on a 250 rpm shaker platform at 37 ^°^C for 18 hrs. Then, the *E. coli* cell population density was counted with a cytometer and the cells were harvested from the liquid culture by centrifugation at 5000 rcf for 10 min. The supernatant from the centrifugation was decanted and the pelleted cells were retained. The pelleted cells were then resuspended in 40 ml of liquid medium containing 20% LB and 80% BG-11 to mimic soil environmental water so that each 40 ml contained 3.33 x 10^11^ recipient cells (8.33×10^9^ cells/mL). An aliquot of 5 ml of the cell suspension (8.33×10^9^ cells/mL) was placed into one of the six 10-ml rectangular flasks in the presence or absence of 1 µg of pAM5409 DNA for the HGT liquid incubation assay that lasted up to 5 days. Following 1, 3 and 5 days of liquid incubation, the incubated recipient cell cultures were sampled for plating (100 µl per plate) onto six solid LB medium plates in the presence of antibiotics (Sp/Sm) for colony formation unit assay or in the absence of antibiotics as a control. The number of colony formation unit were counted manually and digitally using OpenCFU and FIJI for ImageJ. The experimental results demonstrated that HGT could indeed occur through direct uptake of plasmid DNA from the external liquid phase as evidenced by the expression of the yellow fluorescent protein reporter in colonies that were observed with fluorescent microscopic techniques. However, the frequency of HGT through direct uptake of DNA from the external liquid phase appeared to be quite low, which is just slightly higher than the frequency (rate) of bacterial spontaneous natural mutations. It requires a carful effort to detect and analyze this mode of HGT as described with more details as follows.

**Figure 2.**
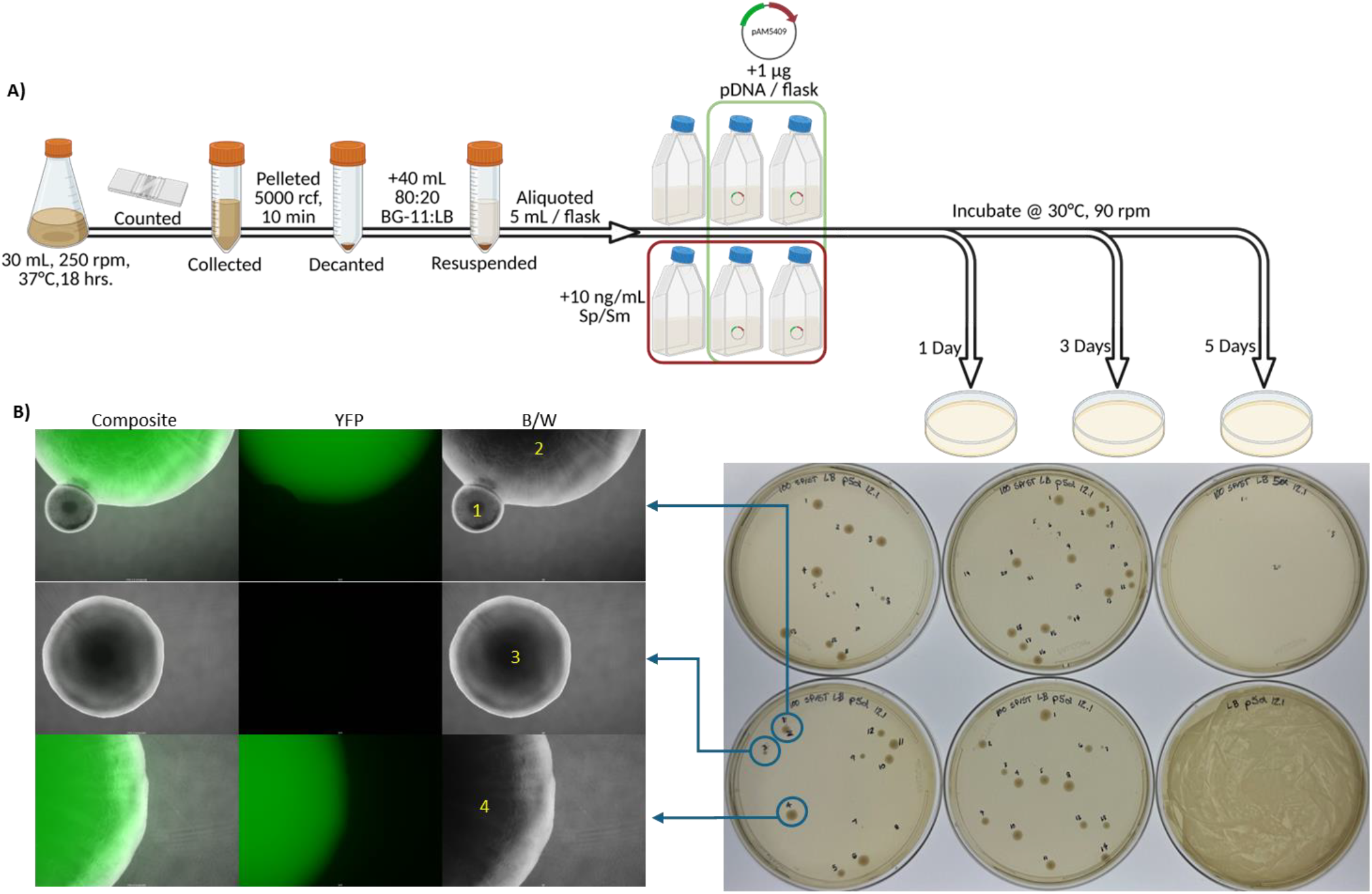
Experimental design and data acquisition. A) Illustration depicting the experimental flow & sampling schedule implemented for K12 strain of *E. coli*; process was identical for DH5α and 10B. B) Plate CFUs (right) were either manually counted (as depicted) or cropped down to individual plate images and counted with software when colonies numbered in the hundreds. Microscopic images (left) of manually counted colonies were taken in polarized B/W and using a YFP filter cube (**Figure S3**). Colonies exhibiting mean YFP intensity greater than 10 times that of mean background YFP intensity were considered YFP expressors for probability calculations.

Under all experimental conditions evaluated, positive control agar plates (without any antibiotic) exhibited unrestrained bacterial growth resulting in a “lawn” of colonies for all 3 *E. coli* strains, indicating that all nutrient-limited liquid incubation recipient cell suspensions contained ample viable bacterial cells for the experiment duration irrespective of sub-lethal Sp/Sm (10 ng/ml) exposure (**Figures S1 and S2**). Despite this, positive control growth was the only observed K12 growth; all other K12 plates (containing antibiotic Sp/Sm 100 µg /ml) failed to develop any visible colonies. This is not surprising considering that K12 has fully functional *mcr, mrr* and *hsd* restriction modification systems. The remaining two strains, DH5α and DH10B, both exhibited natural competence as evidenced by the formation of YFP expressing colonies as well as acquired antibiotic Sp/Sm resistance as evidenced by non-YFP expressing colonies. Interestingly, our negative controls for the DH5α and 10B strains ([Sp/Sm] 100 μg/mL agar inoculated with 100 µl of liquid incubated recipient cell culture without pAM5409) developed a number of non-YFP expressing CFUs comparable to the number observed on plates originating from the incubation liquid containing pAM5409 and recipient cells. This result indicated that the frequency of HGT through direct uptake of DNA from the external liquid phase was quite low, which is just slightly higher than the frequency (rate) of bacterial spontaneous natural mutations.

Both DH5α and 10B consistently formed YFP expressing colonies across all replicate plates after 1 day of liquid incubation with the eDNA (pAM5409). From [Sp/Sm] 0 ng/mL liquid incubation of recipient cells with pAM5409, both DH5α and 10B formed a mean number of 10 YFP CFUs per plate; from [Sp/Sm] 10 ng/mL liquid incubation of recipient cells with pAM5409, DH5α formed a mean of 11 YFP CFUs per plate while 10B formed a mean of 22 YFP CFUs per plate. Additionally, [Sp/Sm] 0 ng/mL 10B cultures formed a mean of 4 YFP CFUs over 3 replicates after 3 days of liquid incubation. In addition to forming YFP expressing colonies, both DH5α and 10B also formed non-YFP expressing CFUs seemingly independent of pAM5409 presence or sub-lethal Sp/Sm exposure. After 1, 3 and 5 days of liquid culturing, DH5α from [Sp/Sm] 0 ng/mL liquid incubation formed a mean of 5, 420, and 134 CFUs per antibiotic plate while [Sp/Sm] 10 ng/mL liquid DH5α incubation yielded 8, 87, and 169 mean CFUs per antibiotic plate, respectively. Conversely, [Sp/Sm] 0 ng/mL liquid 10B incubation formed a mean of 4, 1, and 0 CFUs per plate after 1, 3, and 5 days of liquid incubation, while [Sp/Sm] 10 ng/mL liquid 10B incubation yielded 2, 2, and 1 mean CFUs per plate (**Figure 3**).

**Figure 3.**
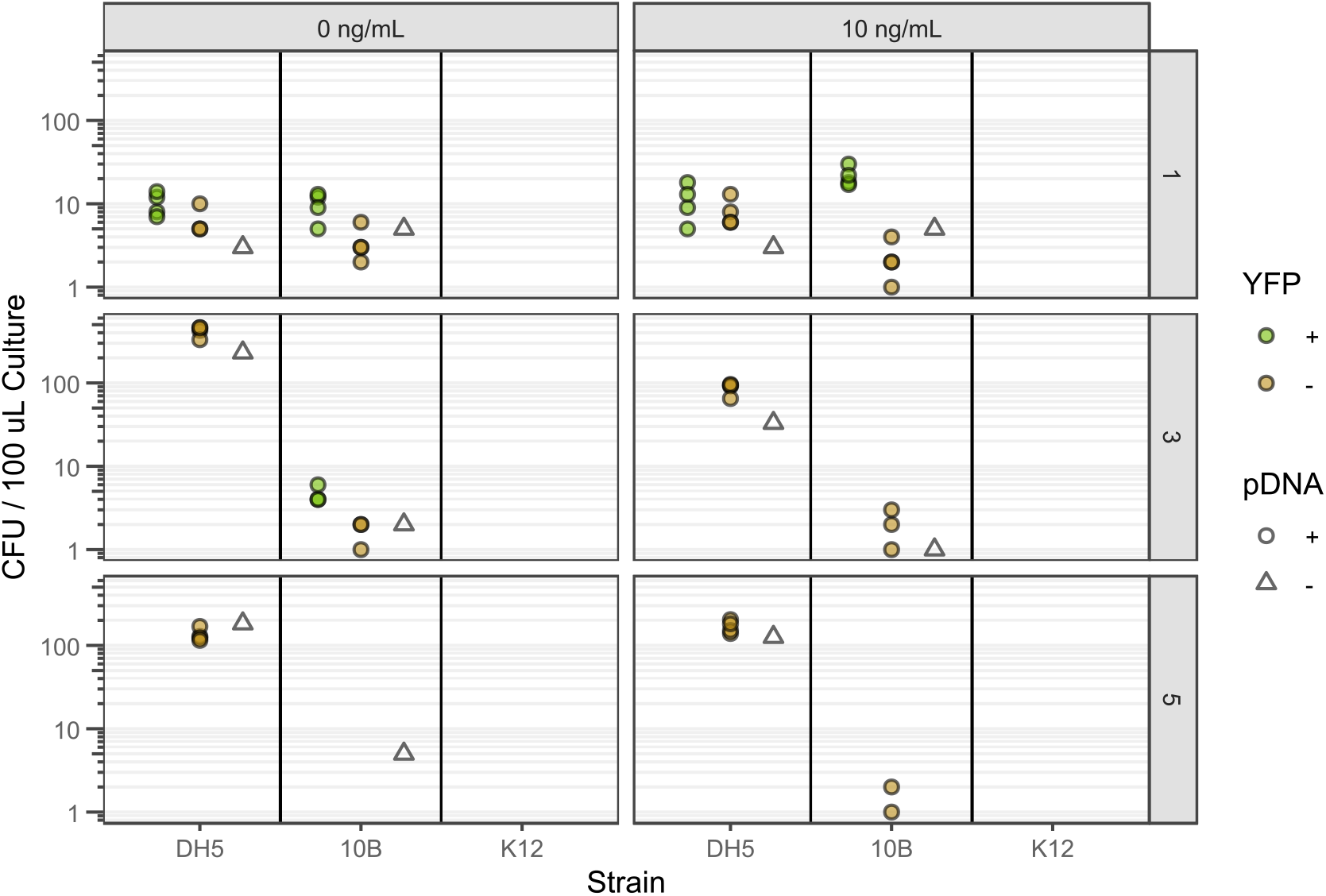
Plots of colony forming units (CFU) per 100 μL of incubated liquid recipient cells suspension. Panel columns denote CFUs from liquid cultures without and with Sp/Sm (0 or 10 ng/mL) while panel rows represent CFUs obtained from liquid recipient cells suspensions incubated for the indicated number of days (1, 3, 5). Within each individual plot, points represent the number of CFUs per plate observed colored by the absence of (brown) or the expression of (green) YFP. Circles represent CFUs originating from liquid recipient cells suspensions containing pAM5409 (pDNA^+^) while triangles represent CFUs obtained from liquid recipient cells suspensions without pAM5409 (pDNA^-^). CFU axis is log_10_ to permit visualization of all data.

**Figure 4.**
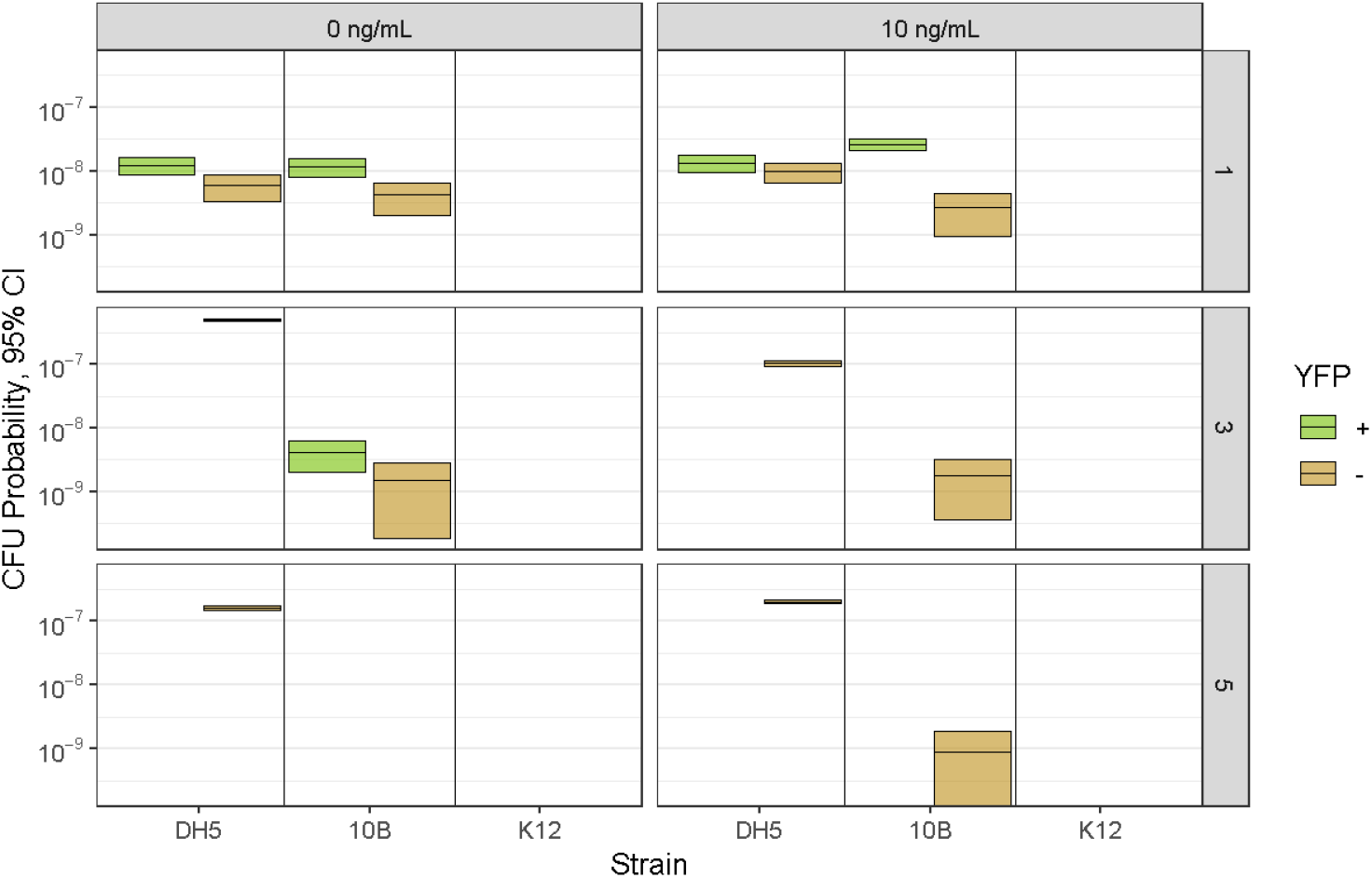
Mean probability of CFU formation and 95% confidence intervals (CI) for *E. coli* strains under tested experimental conditions. Values plotted are those presented in **Table 2**.

Using the observed CFUs and the initial cell density of the liquid cultures, we calculated the probability of 3 types of events: transformation (HGT) with pAM5409 as indicated by the expression of *yfp*, antibiotic resistance without *yfp* expression as indicated by non-YFP colonies that were exposed to pAM5409 and potentially natural mutational antibiotic resistance as indicated by colony formation without exposure to pAM5409 (**Table 2**). Our results indicate that the probability for transformation (HGT) is the highest within the first 24 hours of liquid culturing (10^-8^ vs 10^-9^ after 24 and 72 hours of liquid culturing, respectively) and that DH10B is marginally more competent than DH5α (4.2 x 10^-9^ vs 0 after 3 days of liquid culturing). The presence of 10 ng/mL Sp/Sm had an insignificant effect on the probability of transformation.

**Table 2.** Calculated probability of HGT (transformation), Sp/Sm resistance acquired from either the plasmid or spontaneous natural mutation, and Sp/Sm resistance through spontaneous natural mutation for observed CFUs of each *E. coli* strain after incremental days of liquid incubation with 0 or 10 ng/mL Sp/Sm in 20% LB:80% BG-11 liquid medium.

| Transformation (HGT) Probability: YFP Expressing CFU / Recipient Cell Seeded |  |  |  |  |  |  |
| --- | --- | --- | --- | --- | --- | --- |
| [Sp/Sm], Culture | 0 ng/mL |  |  | 10 ng/mL |  |  |
| Days Cultured | 1 | 3 | 5 | 1 | 3 | 5 |
| K12 | - | - | - | - | - | - |
| DH5α | $1.2 \times 10^{-8}$<br>$\pm 3.8 \times 10^{-9}$ | - | - | $1.4 \times 10^{-8}$<br>$\pm 6.7 \times 10^{-9}$ | - | - |
| DH10B | $1.2 \times 10^{-8}$<br>$\pm 4.3 \times 10^{-9}$ | $4.2 \times 10^{-9}$<br>$\pm 3.0 \times 10^{-9}$ | - | $2.6 \times 10^{-8}$<br>$\pm 7.1 \times 10^{-9}$ | - | - |
| Transformation and/or Spontaneous Mutation: Sp/Sm Resistant, Non-YFP Expressing CFU in the Presence of eDNA / Recipient Cell Seeded |  |  |  |  |  |  |
| K12 | - | - | - | - | - | - |
| DH5α | $6.0 \times 10^{-9}$<br>$\pm 4.9 \times 10^{-9}$ | $5.04 \times 10^{-7}$<br>$\pm 7.41 \times 10^{-8}$ | $1.61 \times 10^{-7}$<br>$\pm 2.90 \times 10^{-8}$ | $9.9 \times 10^{-9}$<br>$\pm 4.0 \times 10^{-9}$ | $1.0 \times 10^{-7}$<br>$\pm 1.8 \times 10^{-8}$ | $2.03 \times 10^{-7}$<br>$\pm 3.55 \times 10^{-8}$ |
| DH10B | $4.2 \times 10^{-9}$<br>$\pm 2.1 \times 10^{-9}$ | $1.5 \times 10^{-9}$<br>$\pm 1.2 \times 10^{-9}$ | - | $2.7 \times 10^{-9}$<br>$\pm 1.5 \times 10^{-9}$ | $1.8 \times 10^{-9}$<br>$\pm 1.6 \times 10^{-9}$ | $9.0 \times 10^{-10}$<br>$\pm 1.2 \times 10^{-9}$ |
| Spontaneous Mutation: Sp/Sm Resistant CFU in the Absence of eDNA / Recipient Cell Seeded |  |  |  |  |  |  |
| K12 | - | - | - | - | - | - |
| DH5α | $3.6 \times 10^{-9}$ | $2.77 \times 10^{-7}$ | $2.20 \times 10^{-7}$ | $3.6 \times 10^{-9}$ | $4.0 \times 10^{-8}$ | $1.51 \times 10^{-7}$ |
| DH10B | $6.0 \times 10^{-9}$ | $2.4 \times 10^{-9}$ | - | $6.0 \times 10^{-9}$ | $1.2 \times 10^{-9}$ | - |

### *E. coli* growth assay over 48 hours as a function of percent LB in medium

Lysogeny broth (LB) medium formulations (which comprise organic nutrients peptides and casein peptones, vitamins, plus minerals and trace elements) have been an industry standard for the growth of bacteria including *Escherichia coli* (*E. coli*) ^30^. BG-11 media (which comprise only inorganic nutrients including major minerals P, N, K, Na, Ca, Mg and trace elements) can support photoautotrophic growth of freshwater photosynthetic microorganisms such as blue-green algae ^31^. Therefore, we considered a mixture of LB and BG-11 to mimic soil-environmental water for the HGT assays with pAM5409 as eDNA.

To better understand the effect of incubation liquid nutrient content on recipient cell growth and maintenance, *E. coli* strains K12, DH5α and DH10B were incubated in different mediums containing increasing fractions of LB mixed with BG-11 over an LB range of 0% to 100% (**Figure 5**). With the exception of 0% LB (100% BG-11) cultures, all 3 strains exhibited an initial rapid increase in cells population density associated with the exponential (logarithmic) growth phase for the first 8-20 hrs followed by a stationary growth phase in 5% LB (95% BG-11), 10% LB (90% BG-11), 15% LB (85% BG-11), 20% LB (80% BG-11), 25% LB (75% BG-11), 30% LB (70% BG-11), 40% LB (60% BG-11), 50% LB (50% BG-11), and 100% LB (0% BG-11). Across all strains, as the % LB was increased the gradual decline in cell population density characteristic of stationary phase growth (5-30% LB) was reversed to a gradual increase in cell density (40-100% LB). All strains transitioned from exponential to stationary phase within the first 8 to 20 hrs with a general trend of later transitioning corresponding to cultures containing a greater percentage of LB in the medium. From these results, we selected an organic nutrient-limited liquid medium comprising 20% LB and 80% BG-11 to support both sufficient growth and sustained maintenance of *E. coli*.

**Figure 5.**
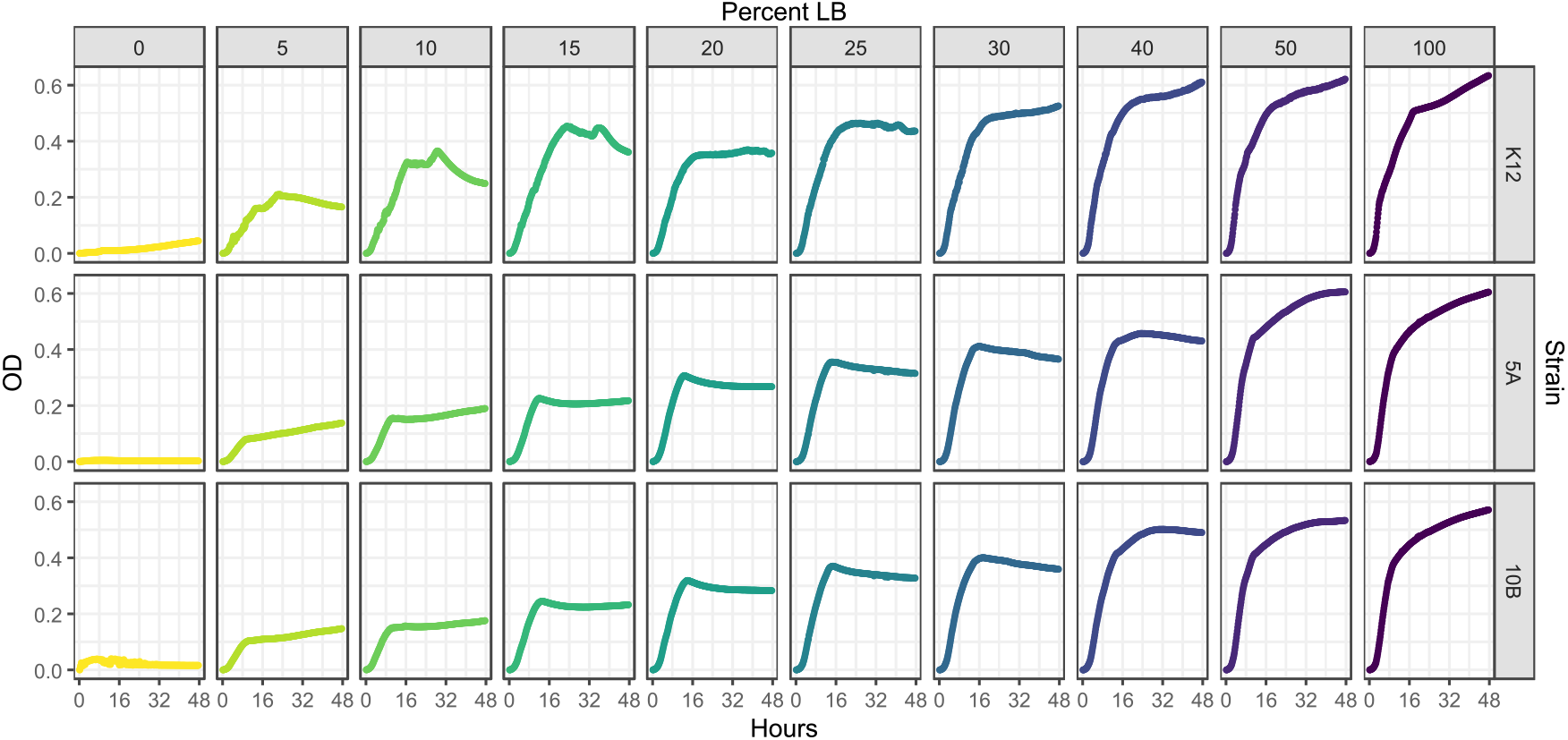
Optical density (OD) measured every 10 minutes for 48 hours of K12, DH5α (5A) & DH10B (10B) in BG-11 medium containing an increasing percent of LB. OD is defined as the absorbance of wavelength 600 nm light. Cultures were run in parallel with blanks of corresponding identical medium composition. Plotted values are the mean of 2 liquid culture replicates minus the mean of 2 corresponding blanks at every time point.

## Discussion and conclusion

The experimental results seems to indicate that DH5α and 10B are naturally competent during the exponential growth phase within the first day in an organic-nutrient-limited liquid culture with a probability of 1.2 – 1.4 x 10^-8^ CFU/seeded recipient cell for DH5α and 0.43 – 2.6 x 10^-8^ CFU/seeded recipient cell for DH10B. This contrasts with previously held beliefs that *E. coli* is not naturally competent, although more recent findings are calling that belief into question^13,32^. A thorough 2018 review by Hasegawa, Suzuki & Maeda of *E. coli* competency reported in literature since 2000 found natural transformation frequencies under a plethora of different conditions. Of relevance are similar transformation frequences (10^-6^ – 10^-9^) obtained in experiments examining the effects of different food stuffs and solid-air biofilms on *E. coli* acquisition of supplied plasmid ^32^. It is well established that the efficiency of genetic transformation increases in the presence of divalent cations ^33^; our nutrient-limited liquid medium has a divalent cation concentration of 0.45 mM, of which 0.44 mM is Mg^2+^ and Ca^2+^, resulting from its 80% BG-11 composition and is similar to that found in some of the foods evaluated in the experiments reviewed by Hasegawa, Suzuki & Maeda ^32^. Therefore, the concentration of divalent cations used in our experimental conditions could be a factor contributing to the observed probability of HGT observed in samples from the liquid incubation assays.

Other studies^19^ have also observed diminishing transformation rates as the duration of liquid culturing increases. This trend can be explained by the metabolic transition that occurs when a culture exits exponential phase growth and enters stationary phase. Limited nutrients induce a starvation response which downregulates the σ^70^ RNA polymerase subunit driving the transcription of a host of cellular proliferation genes while upregulating the σ^56^ subunit that reduces cellular division, protein production and non-essential ATP driven processes such as DNA uptake via T4SS import machinery. In the experiments, we observed that while neither DH5α nor DH10B colonies exhibited YFP fluorescence after 5 days of the liquid incubation assay, the number of Sp/Sm resistant yet non-YFP expressing DH5α colonies increased nearly 100 fold after 3 days of the liquid incubation assay independent of sub-lethal Sp/Sm concentration or exposure to the plasmid gene conferring antibiotic resistance. It is assumed that these colonies developed antibiotic resistance either via chromosomal incorporation of the *aadA* gene from the supplied plasmid or via a spontaneous natural mutation in the case of colonies sourced from the liquid culture never exposed to the plasmid pAM5409. For plates from days 3 and 5 of the liquid incubation assay, the number of non-YFP expressing CFUs sourced from liquid recipient cell incubation without pAM5409 were consistent with the number of CFUs sourced from liquid recipient cell incubation containing pAM5409 suggesting a similar mechanism for acquiring antibiotic resistance independent of the supplied eDNA. Since both DH5α and DH10B have loss of function *recA1* mutations which reduce the rate of plasmid recombination, the latter option seems more likely.

It was qualitatively observed that the radius of non-YFP expressing colonies were consistently smaller than that of YFP expressing colonies implying attenuated growth. While this observation is consistent with DH5α’s inability to synthesize thiamine due to the *thi-1* mutation, it contradicts the 100 fold CFU increase exhibited by DH5α after 3 and 5 days of liquid incubation assay when medium thiamine was certainly depleted given the duration of the incubation. Our growth assays (Figure 5) indicate that all 3 strains reach stationary phase after 12-15 hours independent of the amount of amino acid concentration supplied by the LB; if thiamine supply were attenuating the growth of DH5α, one would expect a reduction in the number of CFUs observed, not an increase. Given that Sp/Sm inhibits protein synthesis via ribosomal 30S binding, it is also plausible that point mutations in the *rps* gene could result in an attenuated antibiotic efficiency while still allowing sufficient protein synthesis to enable cell survival. Regardless, given the lack of knowledge surrounding the induction of bacterial “persisters,” antibiotic tolerant bacteria that arise from starvation conditions, the increase in CFUs after multiple days of liquid incubation may be the result of yet unidentified interplay between bacterial survival mechanisms ^34^.

The horizontal gene transfer (HGT) frequency may be calculated as the HGT probability per cell per time unit (day) using Eq. 2 as described in the method section. Therefore, based on the YFP expressing CFU per recipient cell observed within the 1 day liquid incubation period, the HGT frequency was calculated to 1.2 x 10^-8^ per cell day for both DH5α and DH10B. Similarly, the frequency of spontaneous natural mutation for natural Sp/Sm resistance development was estimated to 4.8 10^-9^ per cell as calculated from 3.6 x 10^-9^ per cell day for DH5α and 6.0 x 10^-9^ per cell day for DH10B.

As shown in Figure 5, the result of *E. coli* growth assay as a function of % LB in liquid medium showed that the exponential (logarithmic) cells population growth phase occurs within the first day (in the first 8 to 20 hours); after that (in days 2 to 5) was the stationary population phase. Therefore, based on the HGT assay result (Table 2) that the extension of the liquid incubation of plasmid DNA and recipient cells to 3 and 5 days did not substantially increase the occurrence of HGT, it appears that the logarithmic cells population growth phase may play a substantial role for HGT through direct eDNA (pAM5409) uptake from the extracellular liquid phase, under the current experimental conditions with nutrient-limited liquid co-incubation medium (20% LB + 80 BG-11).

In conclusion, events of HGT through direct uptake of eDNA were experimentally demonstrated. This experimental study showed that the HGT frequency by direct uptake of plasmid DNA may be 1.2 x 10^-8^ per cell day for both DH5α and DH10B. The extension of the liquid incubation of plasmid DNA and recipient cells from 1 day to 3 and 5 days did not substantially increase the HGT frequency. The logarithmic cells population growth phase within the first day (in the first 8 to 20 hours) may play a substantial role for HGT through direct eDNA (pAM5409) uptake from the extracellular liquid phase under the current experimental conditions with nutrient-limited incubation medium (20% LB + 80 BG-11). Under the current experimental conditions, HGT in *E. coli* strains K12 which has its intact genes encoding restriction enzymes and endonucleases (Table 1) seemed to be undetectable. This indicates that K12’s restriction enzymes and endonucleases are likely to play a substantial role to degrade foreign DNA like plasmid pAM5409. Even if K12 uptakes plasmid pAM5409 like its derivatives DH5α and DH10B do, its frequency to retain and express pAM5409 could be so low that its HGT could be “undetectable” under the current experimental conditions. Considering the enormous scale of the environment, even transformation frequencies of 10^−9^−10^−10^ cannot be underestimated as they will have an impact on microbial populations ^32^. Further investigation into HGT of other environmental microorganisms such as other soil bacteria other than *E. coli* should be encouraged. Meanwhile, further investigation into HGT mechanisms such as the one that may underly the antibiotic resistance developed by non-YFP expressing colonies is also warranted given the permissive spread of antibiotic resistant strains in natural microbial ecosystems.

## Methods and Materials

### E. Coli growth assay

One mL of LB in microcentrifuge tubes were inoculated with *E. coli* K12, DH5α & DH10B strains (ThermoFisher Scientific) and cultured overnight at 37° C. The following day a 96 well plate was prepared with 8 sets of wells; each set contained wells with the following ratios of LB:BG-11 medium: 0:100, 5:95, 10:90, 15:85, 20:80, 25:75, 30:70, 40:60, 50:50 and 100:0. All wells had a final volume of 200 μL. The overnight cultures were examined microscopically to ensure viability before inoculating the 96 well plate. Two sets of wells were inoculated per bacterial strain with 5 μL of culture per well; 2 sets of wells were not inoculated and served as blanks. The 96 well plate was covered and incubated at 30°C with intermittent shaking at 60 rpm in a Varioskan Lux multiplate reader for 48 hours with the optical density at a wavelength of 600 nm recorded every 10 min. Replicate absorbances were averaged and the mean blank absorbance was subtracted from the mean culture absorbances at each time point in MS Excel.

### Preparation of plasmid pAM5409

DH5αMCR containing pAM5409 was cultured in 200 mL of LB overnight at 37°C with agitation. Cells were collected via centrifugal pelleting at 5000 rcf for 10 min. Plasmid pAM5409 was extracted and purified using a Nucleobond Xtra Midi kit (Macherey-Nagel) per the manufacturer’s instructions; eluate was further cleaned using a Monarch Spin PCR & DNA Cleanup Kit (NEB) and concentrated by eluting with 500 μL 10mM Tris. Plasmid purity and concentration were determined via triplicate NanoDrop reads and confirmed by PCR amplification and gel electrophoresis of the 2095nt region containing genes *aadA & yfp*. Antibiotic stocks of Spectinomycin dihydrochloride pentahydrate (Sigma Aldrich) and Streptomycin sulfate (Sigma Aldrich) were prepared in nuclease-free water, sterile filtered and stored at 0°C. Solid media plates were prepared by combining equal volumes of autoclaved 2x LB and autoclaved 3% w/w agar in dH_2_O, supplementing with antibiotics as needed to a final concentration of 100 ug/mL and dispensing 25 mL aliquots into disposable petri dishes. Once set, plates were stored at 4°C until use. All solid media plates were spread with 100 uL of inoculum using sterile glass beads and incubated at 37°C for 48 hours before counting colonies and imaging.

### Liquid incubation of *E. coli* cells with pAM5409

*E. coli* K12, DH5α & DH10B strains (ThermoFisher Scientific) were cultured overnight in 30 mL of LB overnight at 37°C with agitation; cell densities were calculated from visual counts using a Petroff-Hausser Counter (VWR, #15170-048) and pelleted via centrifuging at 5000 rcf for 10 min. The pellet was retained by removing the supernatant and the pelleted cells were resuspended in 40 mL of an organic-nutrient-limited liquid medium comprised 80% BG-11 and 20% LB. To seed all replicate cultures with an equivalent number of cells, the cultured strain containing the least number of cells (DH5α, 3.33 x 10^11^ cells) was used as the limiting cell culture stock. Resuspended K12 and 10B had 7.41 mL and 6.49 mL respectively removed and replaced with organic-nutrient-limited liquid medium (80% BG-11 and 20% LB), generating 40 mL of each bacterial strain liquid culture, each containing 3.33 x 10^11^ cells. For each *E. coli* strain, 5 mL of liquid cell culture was aliquoted into one of six 10 mL, untreated, vented Corning flasks (Fisher Scientific, #08-757-500). Flasks were divided into 2 sets of 3 each; one set received no antibiotic (AB^-^) and the other was brought to a sub-lethal [Sp/Sm] concentration of 10 ng/mL (AB^+^). Within each set of 3 flasks, 2 flasks received 1 ug of pAM5409 DNA. All flasks were then incubated at 30°C, 90 rpm in an IKA KS3000i orbital shaking incubator for a total of 5 days (**Figure 3**).

### Plating & imaging

After each of 1, 3 and 5 days of liquid incubation assays, the incubated liquid recipient cells suspensions were sampled and a total of 6 LB agar plates were spread from each set of K12, DH5α & 10B incubation culture flasks: 2 antibiotic (100 µg /ml Sp/Sm) plates from each flask containing plasmid, one antibiotic (100 ug/mL Sp/Sm) plate from each flask without plasmid (negative control for antibiotic selection) and one antibiotic-free plate from a flask containing plasmid (positive control for media growth). All LB agar plates were incubated at 37^°^C for 2 days before being photographed. CFUs/plate were determined either manually or digitally using OpenCFU and FIJI for ImageJ. YFP expression was evaluated microscopically and imaging of colonies was performed on an Olympus BX-43 with a DP-72 camera under magnification with a 10x/0.25 Plan objective. An X-Cite 120Q equipped with an OSRAM HXP-R120W/45C mercury arc lamp provided collimated excitation light via a liquid light guide through a Chroma #49003 filter set (Ex:500/20, 515LP, Em:525/50) for evaluating YFP expression. All images were taken using CellSense software with 50 ms exposures (ISO200) at 10 bit depth and 4096 x 3096 px. resolution.

### Calculation of event probability and frequencies

Event probability presented in **Table 2** were calculated assuming each CFU originated from a single cell with a binomial probability *p* of survival. Dividing the total CFUs across 4 replicate plates by the initial cell population density of the liquid culture multiply with the volume of liquid culture plated (8.33×10^6^ cells/μL x 100 μL/plate x 4 plates) yields the survival probability, an estimate of *p*. Margins of error (±*E*) are reported at the 95% confidence interval (*z*_95%_ = 1.96) by the following equation where *p* is the CFU probability and *n* is the theoretical number of cells plated based on the initial cell population densities:

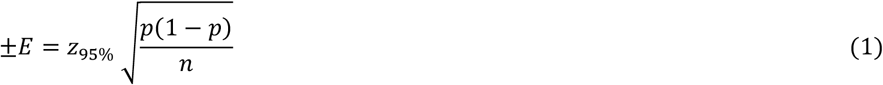

The horizontal gene transfer (HGT) frequency was calculated from the number (*N*_*colony*_) of HGT recipient (*E. coli*) colonies observed per antibiotic LB agar selective plate, the volume (*V*_*Incubation*_) of the donor-recipient co-incubation cell suspension liquid that was used in spreading onto the surface of an antibiotic-containing LB agar medium plate, the concentration (*C*_*Recipient*_) of the initial recipient (*E. coli*) cells in the co-incubation liquid, and the donor-recipient co-incubation time (*t*_*Incubation*_) according to the following HGT Frequency equation ^20^ :

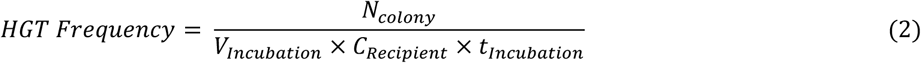

## Acknowledgements

We thank Prof. James W Golden and his associate Bryan Bishé of the University of California at San Diego for their wonderful technical support including helpful discussions and email communications about the pAM5409, which was a gift from Susan Golden (Addgene plasmid # 132662; http://n2t.net/addgene:132662; RRID:Addgene_132662.). We acknowledge the Chem-497 and Chem-498 Undergraduate Student Independent Research Program well prepared Z.S. to participate in this USDA biosafety assessment research project.

## Author contributions statement

J.W.L. and C.W. conceived the experiments. C.W., Z.S. and L.J. conducted experiments and performed imaging. C.W. processed images & data, analyzed results and drafted manuscript. J.W.L. edited and finalized the manuscript. All authors reviewed the manuscript.

## Additional information

The authors declare no competing interests.

## Data availability

All data generated or analyzed during this study are included in this published article and its Supplementary Information available online at the journal website.

## References

1. Brito, I. L. Examining horizontal gene transfer in microbial communities. Nature Reviews Microbiology 19, 442–453, doi:10.1038/s41579-021-00534-7 (2021).

2. van Gestel, J. et al. Short-range quorum sensing controls horizontal gene transfer at micron scale in bacterial communities. Nature Communications 12, 2324, doi:10.1038/s41467-021-22649-4 (2021).

3. Seitz, P. & Blokesch, M. Cues and regulatory pathways involved in natural competence and transformation in pathogenic and environmental Gram-negative bacteria. FEMS Microbiol Rev 37, 336–363, doi:10.1111/j.1574-6976.2012.00353.x (2013).

4. Lang, A. S., Buchan, A. & Burrus, V. Interactions and evolutionary relationships among bacterial mobile genetic elements. Nature Reviews Microbiology, doi:10.1038/s41579-025-01157-y (2025).

5. Borodovich, T., Shkoporov, A. N., Ross, R. P. & Hill, C. Phage-mediated horizontal gene transfer and its implications for the human gut microbiome. Gastroenterology Report 10, doi:10.1093/gastro/goac012 (2022).

6. Álvarez-Rodríguez, I. et al. Type IV Coupling Proteins as Potential Targets to Control the Dissemination of Antibiotic Resistance. Frontiers in Molecular Biosciences 7, doi:10.3389/fmolb.2020.00201 (2020).

7. Costa, T. R. D., Patkowski, J. B., Macé, K., Christie, P. J. & Waksman, G. Structural and functional diversity of type IV secretion systems. Nature Reviews Microbiology 22, 170–185, doi:10.1038/s41579-023-00974-3 (2024).

8. Goldlust, K. et al. The F pilus serves as a conduit for the DNA during conjugation between physically distant bacteria. Proceedings of the National Academy of Sciences 120, e2310842120, doi:10.1073/pnas.2310842120 (2023).

9. Mandel, M. & Higa, A. Calcium-dependent bacteriophage DNA infection. J Mol Biol 53, 159–162, doi:10.1016/0022-2836(70)90051-3 (1970).

10. Vesel, N. et al. Naturally competent bacteria and their genetic parasites—a battle for control over horizontal gene transfer? FEMS Microbiology Reviews 49, doi:10.1093/femsre/fuaf035 (2025).

11. Niu, C. et al. Molecular mechanisms and applications of natural transformation in bacteria. Front Microbiol 16, 1578813, doi:10.3389/fmicb.2025.1578813 (2025).

12. Bergmans, H. E., van Die, I. M. & Hoekstra, W. P. Transformation in Escherichia coli: stages in the process. J Bacteriol 146, 564–570, doi:10.1128/jb.146.2.564-570.1981 (1981).

13. Sinha, S. & Redfield, R. J. Natural DNA uptake by Escherichia coli. PLoS One 7, e35620, doi:10.1371/journal.pone.0035620 (2012).

14. Barnes, M. A. & Turner, C. R. The ecology of environmental DNA and implications for conservation genetics. Conservation Genetics 17, 1–17, doi:10.1007/s10592-015-0775-4 (2016).

15. Kittredge, H. A., Dougherty, K. M. & Evans, S. E. Dead but Not Forgotten: How Extracellular DNA, Moisture, and Space Modulate the Horizontal Transfer of Extracellular Antibiotic Resistance Genes in Soil. Appl Environ Microb 88, doi:ARTN e02280-21 10.1128/aem.02280-21 (2022).

16. Wang, Z. et al. Critical roles of cyanobacteria as reservoir and source for antibiotic resistance genes. Environ Int 144, 106034, doi:10.1016/j.envint.2020.106034 (2020).

17. Gulden, R. H. et al. Quantitation of Transgenic Plant DNA in Leachate Water: Real-Time Polymerase Chain Reaction Analysis. Journal of Agricultural and Food Chemistry 53, 5858–5865, doi:10.1021/jf0504667 (2005).

18. Foucher, A. et al. Persistence of environmental DNA in cultivated soils: implication of this memory effect for reconstructing the dynamics of land use and cover changes. Scientific Reports 10, 10502, doi:10.1038/s41598-020-67452-1 (2020).

19. Macedo, G. et al. Horizontal Gene Transfer of an IncP1 Plasmid to Soil Bacterial Community Introduced by Escherichia coli through Manure Amendment in Soil Microcosms. Environmental Science & Technology 56, 11398–11408, doi:10.1021/acs.est.2c02686 (2022).

20. Lee, J. W. Protocol measuring horizontal gene transfer from algae to non-photosynthetic organisms. Methodsx 6, 1564–1574, doi:10.1016/j.mex.2019.05.022 (2019).

21. Nguyen, T. H. et al. Demonstration of horizontal gene transfer from genetically engineered Thermosynechococcus elongatus BP1 to wild-type E. coli DH5α. Gene 704, 49–58, doi:10.1016/j.gene.2019.03.014 (2019).

22. Lan, E. I. & Liao, J. C. Metabolic engineering of cyanobacteria for 1-butanol production from carbon dioxide. Metab Eng 13, 353–363, doi:10.1016/j.ymben.2011.04.004 (2011).

23. Atsumi, S., Higashide, W. & Liao, J. C. Direct photosynthetic recycling of carbon dioxide to isobutyraldehyde. Nature Biotechnology 27, 1177–1180, doi:10.1038/nbt.1586 (2009).

24. Taton, A. et al. Heterologous Expression in Anabaena of the Columbamide Pathway from the Cyanobacterium Moorena bouillonii and Production of New Analogs. ACS Chemical Biology 17, 1910–1923, doi:10.1021/acschembio.2c00347 (2022).

25. Taton, A. et al. Heterologous Expression of Cryptomaldamide in a Cyanobacterial Host. ACS Synthetic Biology 9, 3364–3376, doi:10.1021/acssynbio.0c00431 (2020).

26. Taton, A. et al. Broad-host-range vector system for synthetic biology and biotechnology in cyanobacteria. Nucleic Acids Res 42, e136, doi:10.1093/nar/gku673 (2014).

27. Bishé, B., Taton, A. & Golden, J. W. Modification of RSF1010-Based Broad-Host-Range Plasmids for Improved Conjugation and Cyanobacterial Bioprospecting. iScience 20, 216–228, doi:10.1016/j.isci.2019.09.002 (2019).

28. Kohanski, M. A., DePristo, M. A. & Collins, J. J. Sublethal antibiotic treatment leads to multidrug resistance via radical-induced mutagenesis. Mol Cell 37, 311–320, doi:10.1016/j.molcel.2010.01.003 (2010).

29. Wistrand-Yuen, E. et al. Evolution of high-level resistance during low-level antibiotic exposure. Nature Communications 9, 1599, doi:10.1038/s41467-018-04059-1 (2018).

30. Sezonov, G., Joseleau-Petit, D. & D’Ari, R. Escherichia coli physiology in Luria-Bertani broth. J Bacteriol 189, 8746–8749, doi:10.1128/jb.01368-07 (2007).

31. Stanier, R. Y., Kunisawa, R., Mandel, M. & Cohen-Bazire, G. Purification and properties of unicellular blue-green algae (order Chroococcales). Bacteriol Rev 35, 171–205, doi:10.1128/br.35.2.171-205.1971 (1971).

32. Hasegawa, H., Suzuki, E. & Maeda, S. Horizontal Plasmid Transfer by Transformation in Escherichia coli: Environmental Factors and Possible Mechanisms. Front Microbiol 9, 2365, doi:10.3389/fmicb.2018.02365 (2018).

33. Asif, A., Mohsin, H., Tanvir, R. & Rehman, Y. Revisiting the Mechanisms Involved in Calcium Chloride Induced Bacterial Transformation. Front Microbiol 8, 2169, doi:10.3389/fmicb.2017.02169 (2017).

34. Wang, M., Chan, E. W. C., Wan, Y., Wong, M.H.-y. & Chen, S. Active maintenance of proton motive force mediates starvation-induced bacterial antibiotic tolerance in Escherichia coli. Communications Biology 4, 1068, doi:10.1038/s42003-021-02612-1 (2021).

