## Supplemental Information for "Horizontal Gene Transfer of *E. coli* by direct DNA uptake from extracellular liquid phase"

for:

Additional experimental data: 3 Figures on pages S2-S6

Figure S1 presents photographs of plates for DH5α, 10B & K12 cultures after 1, 3 & 5 days incubation in reduced LB medium (80% BG-11, 20% LB).

Figure S2 presents photographs of plates for DH5α, 10B & K12 cultures after 1, 3 & 5 days incubation in reduced LB medium (80% BG-11, 20% LB) with 10 ng/mL [Sp/Sm].

Figure S3 presents evidence for the identification of YFP expressing colonies for 2 agar plates.

**Key words**: Horizontal gene transfer, genetic transformation, Bacteria E. coli, direct uptake of DNA from extracellular liquid, biosafety research, environmental DNA

| 0 ng/mL | Day 1 | Day 3 | Day5 |
|---|---|---|---|
| DH5α | 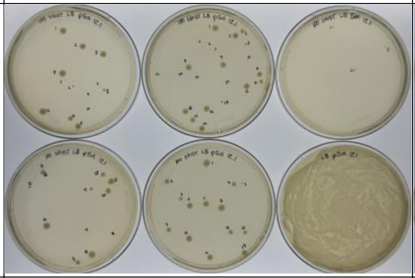 | 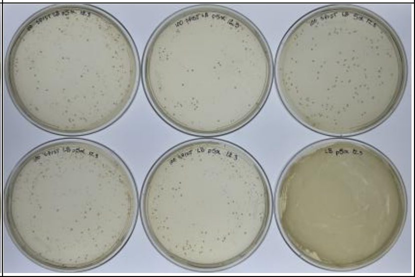 | 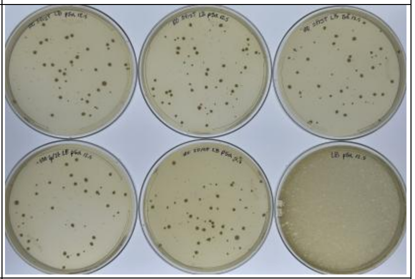 |
| DH10B | 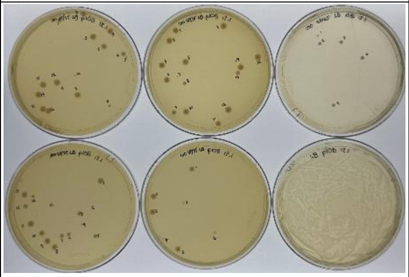 | 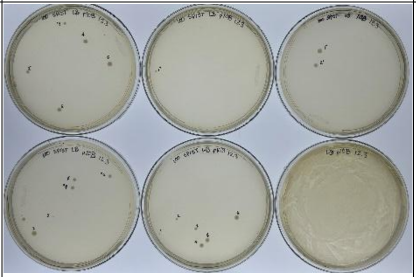 | 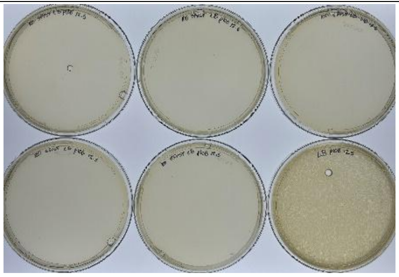 |
| K12 | 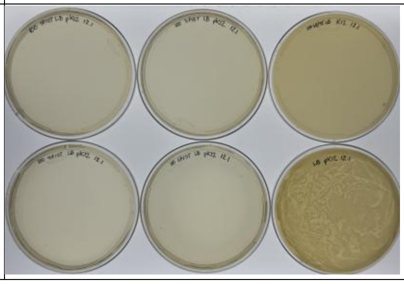 | 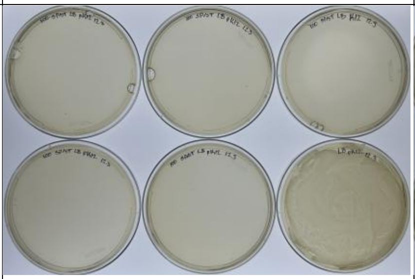 | 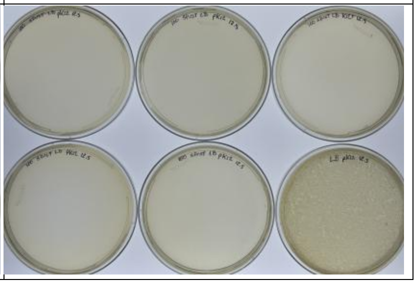 |

**Figure S1.** Image sets of plates for DH5α, 10B & K12 cultures after 1, 3 & 5 days incubation in reduced LB medium (80% BG-11, 20% LB). Each image of 6 plates contains a negative control (upper right), positive control (lower right), 2 plates from 1 replicate culture (upper/lower left) and 2 plates from 2nd replicate culture (upper/lower middle).

| 10 ng/mL | Day 1 | Day 3 | Day5 |
|---|---|---|---|
| DH5α | 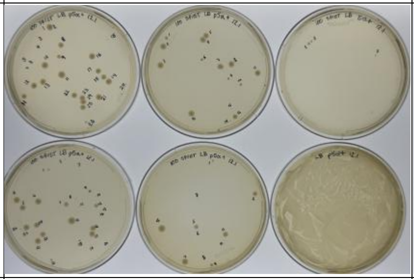 | 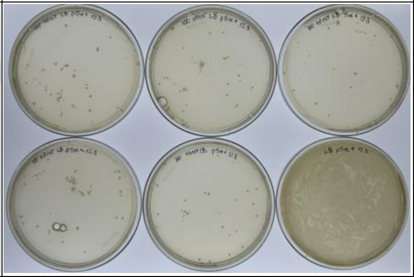 | 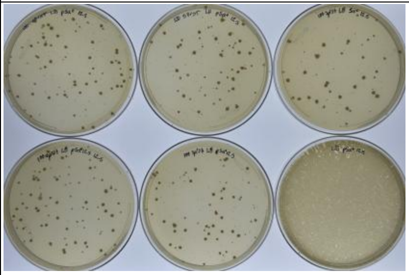 |
| DH10B | 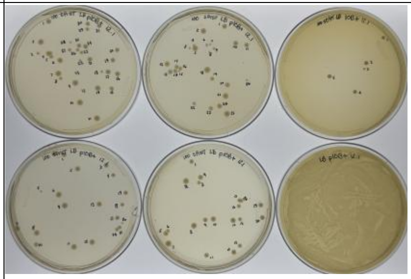 | 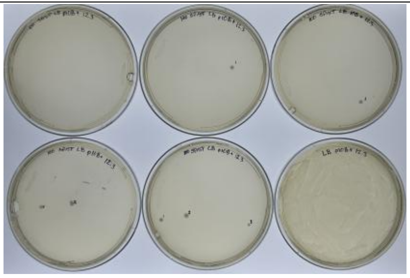 | 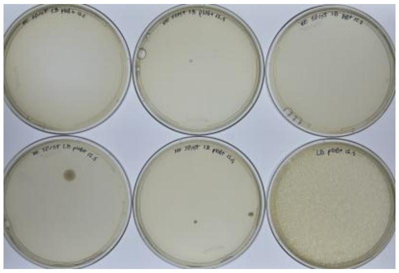 |
| K12 | 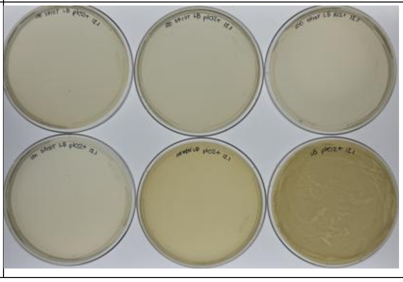 | 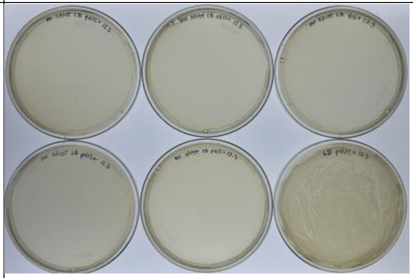 | 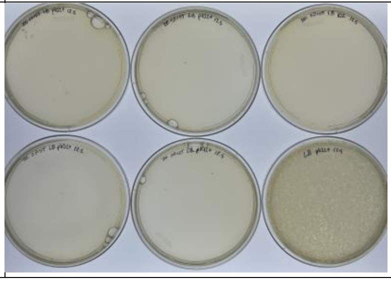 |

**Figure S2.** Image sets of plates for DH5α, 10B & K12 cultures after 1, 3 & 5 days incubation in reduced LB medium (80% BG-11, 20% LB) with 10 ng/mL [Sp/Sm]. Each image of 6 plates contains a negative control (upper right), positive control (lower right), 2 plates from 1 replicate culture (upper/lower left) and 2 plates from 2nd replicate culture (upper/lower middle).

| DH5α.1.1 | DH5α+.1.1 |
| --- | --- |
| 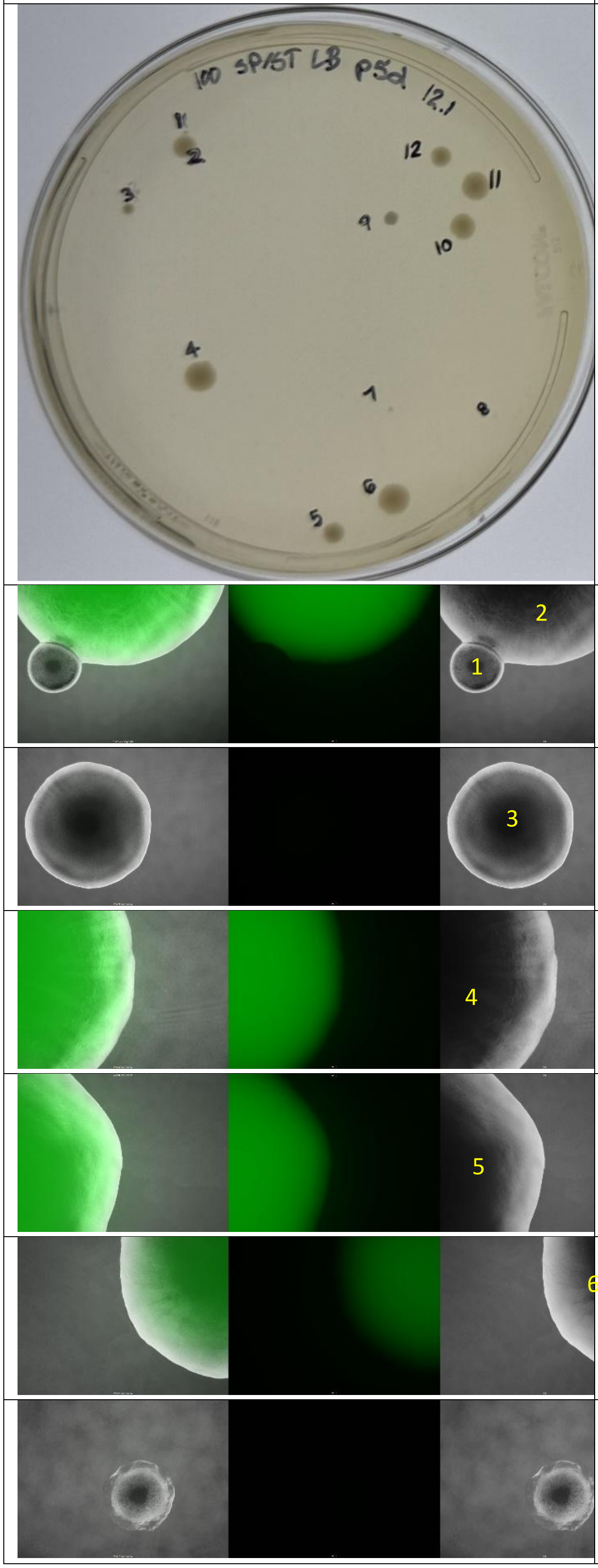 | 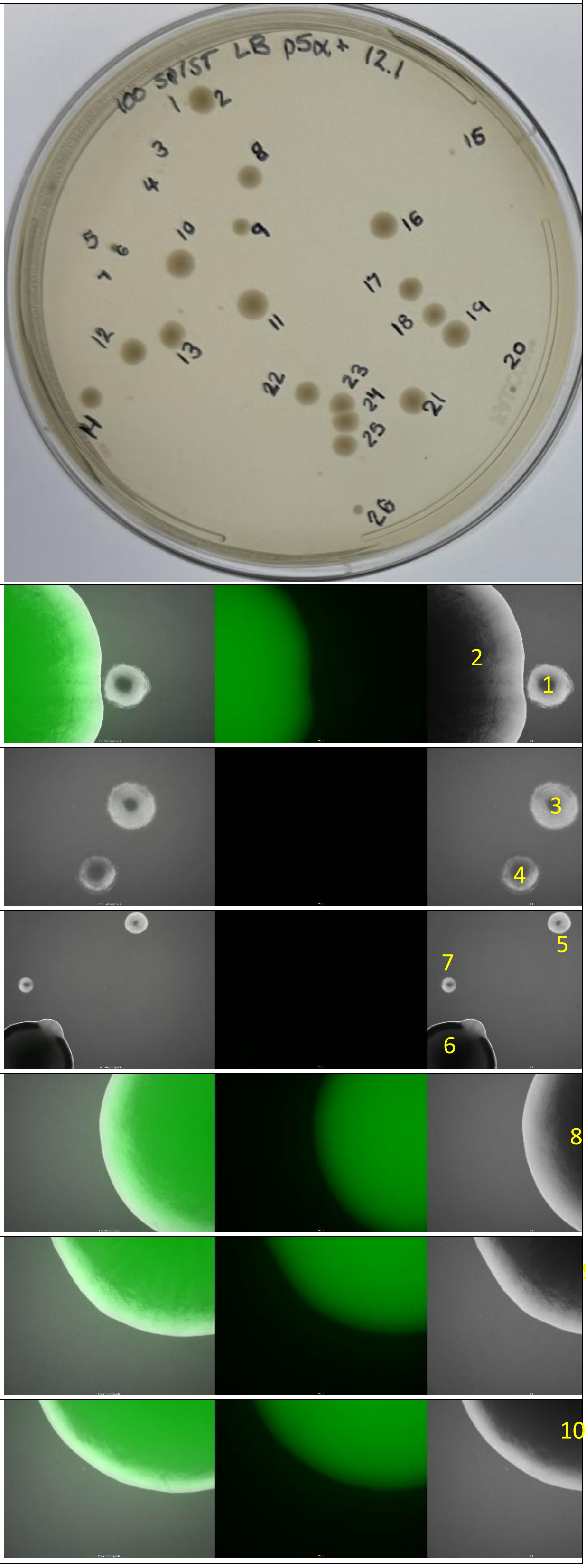 |

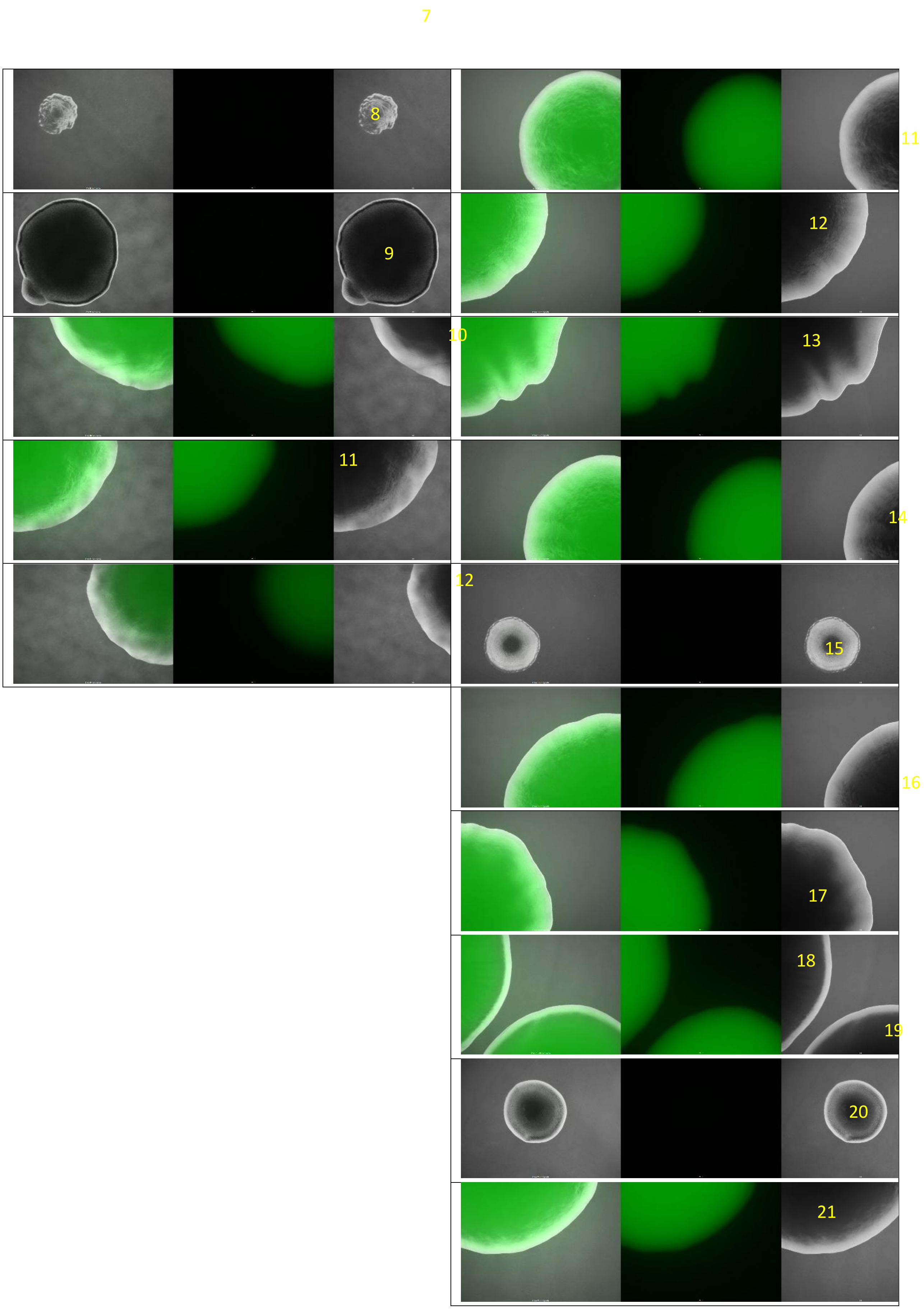
7
8
11
9
12
10
13
11
14
12
15
16
17
18
19
20
21

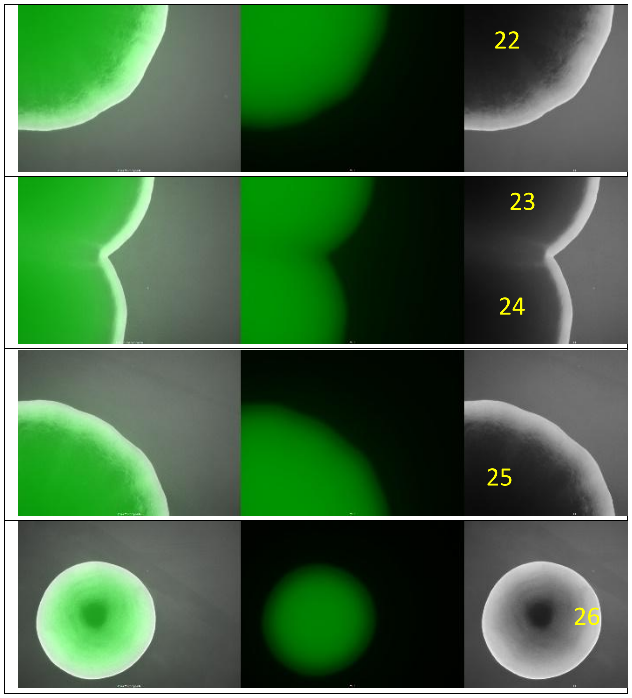


**Figure S3.** Identification of YFP expressing colonies for 2 agar plates. Each column begins with an image of the source plate with colonies manually numbered in black followed by microscopic image panels corresponding to each numbered colony, but in yellow. Each microscopic panel contains a black and white polarized image (right), a YFP filtered image (center) and a composite of the two (left). Panel images were assembled in FIJI for ImageJ using a custom macro for bulk image processing.
